# Long distance homing in the cane toad (*Rhinella marina*) in its native range

**DOI:** 10.1101/2021.06.16.448742

**Authors:** Daniel A. Shaykevich, Andrius Pašukonis, Lauren A. O’Connell

## Abstract

Many animals exhibit complex navigation over different scales and environments. Navigation studies in amphibians have largely focused on species with life histories that require advanced spatial capacities, such as territorial poison frogs and migratory pond-breeding amphibians that show fidelity to mating sites. However, other species have remained relatively understudied, leaving the possibility that well-developed navigational abilities are widespread. Here, we measured short-term space use in non-territorial, non-migratory cane toads (*Rhinella marina*) in their native range in French Guiana. After establishing home range, we tested their ability to return to home following translocations of 500 and 1000 meters. Toads were able to travel in straight trajectories back to home areas, suggesting map-like navigational abilities similar to those observed in amphibians with more complex spatial behavior. These observations break with the current paradigm of amphibian navigation and suggest that navigational abilities may be widely shared among amphibians.

## Résumé

Un grand nombre d’animaux présente une navigation complexe à différentes échelles et dans différents environnements. Les études portant sur la navigation des amphibiens se sont majoritairement intéressées aux espèces dont le cycle de vie nécessite des compétences spatiales avancées, telles que les Dendrobates territoriales, transportant des têtards et les amphibiens migrateurs qui se reproduisent dans les étangs et qui sont fidèles à leurs sites de reproduction. Cependant, d’autres espèces ont été relativement peu étudiées et ces capacités de navigation pourraient être bien plus répandues au sein des amphibiens que nous le pensons. Lors de ces travaux, nous avons quantifié l’utilisation de l’espace d’une espèce de crapaud non territorial et non migrateur (Crapaud buffle, *Rhinella marina*), dans son habitat naturel en Guyane et pendant une courte période de temps. Après avoir établi les limites de leur domaine vital, nous avons testé leur capacité à retourner dans leur site d’origine après les avoir déplacés de 500 et 1000 mètres. Les crapauds étaient capables de retourner vers leur sites de capture de manière directe et rectiligne. Ceci suggère qu’ils possèdent des capacités de navigation analogues à celles observées chez les amphibiens présentant un comportement spatial plus complexe. Ces observations contredisent la théorie actuelle sur la navigation des amphibiens et suggèrent que les capacités de navigation pourraient être plus largement partagées au sein des amphibiens.

## Introduction

Many animals navigate for foraging, migration, or reproduction, from tiny ants to huge whales (Allen, 2013; Wittlinger et al., 2006). Long distance navigation has been extensively studied in species such as sea turtles and migratory birds that offer dramatic examples of large-scale movement (Lohmann et al., 2004; Mouritsen, 2018; Mouritsen and Ritz, 2005). Such studies have been instrumental in uncovering mechanisms, like magnetoreception, that allow for such accurate navigation. However, the navigational abilities of many species that are largely sedentary or that live within relatively small areas are still poorly understood. Even if these animals do not often exhibit notable navigational behaviors, they may still have abilities similar to those studied in organisms known for such abilities. Here, we tested the ability of the cane toad (*Rhinella marina*), a non-territorial, non-migratory nocturnal rainforest amphibian, to return to its home site following translocations exceeding typical movements.

Most studies of amphibian navigation have been carried out in migratory species (Phillips et al., 1995; Sinsch, 2006). For example, the European common toad *Bufo bufo* migrates up to 3 km for breeding and can return to breeding sites in straight trajectories following displacements (Sinsch, 1987). Multiple newt and salamander species can home to breeding sites (Sinsch, 1991); displacement studies of the Californian Red-Bellied newt *Taricha rivularis* showed they can return to natal streams from more than 4 km (Twitty et al., 1964). More recently, Neotropical poison frogs (*Aromobatidae* and *Dendrobatidae*) have been increasingly studied for navigational abilities and spatial cognition on the basis of their complex parental care behaviors. Parent frogs must shuttle tadpoles to deposition sites and return to their territories with high specificity (Liu et al., 2019; Pašukonis et al., 2014a; Pašukonis et al., 2018; Roland and O’Connell, 2015; Stynoski, 2009). In one study, Three-Striped poison frog (*Ameerega trivittata*) males were able to navigate back to home territories following displacements of up to 800 m, exhibiting an ability to return from distances exceeding regular movements (Pašukonis et al., 2018). However, limited effort has gone into studying navigational capabilities in anurans (frogs and toads) that do not exhibit complex spatial behaviors or the need for well developed navigational abilities. With this study, we begin to characterize space use and navigational ability in a non-territorial, non-migratory amphibian species that does not provide parental care and does not have other known life history traits suggestive of highly developed navigation.

Cane toads are large, nocturnal, water breeding amphibians ubiquitous throughout their native range in South and Central America. Females lay thousands of eggs at once in a variety of water sources available in local habitats, ranging from moving creeks and rivers to temporary shallow pools (Evans et al., 1996; Hamilton et al., 2005). Though native to tropical rainforests, cane toads have gained international notoriety as an invasive species, most famously in Australia. Cane toads were introduced in northeast Australia in the mid 1930’s and have continuously spread to encompass over 1 million square kilometers with currently ongoing invasive fronts that have been recorded to expand at up to 50km per year (Urban et al., 2008). The variation in behaviors between native and invasive cane toads has created a dichotomy in the characterization of their spatial behaviors. Native toads and those introduced to areas not environmentally conducive for range expansion are viewed as relatively sedentary and do not perform tasks requiring advanced navigation abilities (Pettit et al., 2016; Ward-Fear et al., 2016; Zug and Zug, 1979). Toads on the invasive front in Australia are considered nomadic or dispersive (Brown et al., 2014; Schwarzkopf and Alford, 2002), and while they may move large distances, they seem not to move towards a specific goal location.

There have been numerous studies examining how and when invasive Australian cane toads move and the physiological characteristics affecting their use of space (Brown et al., 2006; Kelehear and Shine, 2020; Phillips and Shine, 2005; Phillips et al., 2007). Though the spatial ecology of invading toads differs from that of native range toads, general activity patterns are consistent between populations (DeVore et al., 2021). However, despite a few reports on homing in cane toads, such as that of cane toads returning to the same light sources to feed (Brattstrom, 1962) and translocated toads returning to bird ground-nests after being removed (Boland, 2004), no work has systematically quantified the navigational ability of the cane toad using tracking methods. To characterize general space use and determine whether cane toads are capable of long range navigation to a specific goal, we tracked toads and carried out translocation-homing experiments in their native rainforest habitat.

## Methods

### Animals

The experiment was carried out around the Saut Pararé camp (4°02’ N - 52°41’W) of the Nouragues Ecological Research Station in the Nature Reserve Les Nouragues, French Guiana from January to March 2020 (the experiment was terminated early by evacuation due to the COVID-19 pandemic). The area is largely composed of primary lowland rainforest and is bordered by the Arataï river to the south.

Toads were mostly found by visual search at night, but were also located during the day in some instances. Upon capture, toads were photographed, measured snout to vent (SVL), and weighed with a hanging scale (Basetech HS-51, Basetech, Winnipeg, MB, Canada). Animals were captured and tagged (Figure 1) over a transect 300 meters long in close proximity to the Arataï river encompassing ∼9300 m^2^. Individuals were identified by dorsal coloration and wart patterns. A total of 5 females and 9 males were tagged over the course of the experiment. Sex was determined by size, male release calls, and recorded instances of amplexus. Females weighed an average of 1.44 kg (0.79-1.93 kg) and had an average SVL of 21.8 cm (18.0-24.0 cm); males had an average mass of 0.58 kg (0.49-0.65 kg) and an average SVL of 15.0 cm (11.5-18.0 cm). Differences between males and females were significant for SVL (t-test, p < 0.05) and mass (independent 2-group Mann-Whitney U Test, p < 0.05).

**Figure 1.**
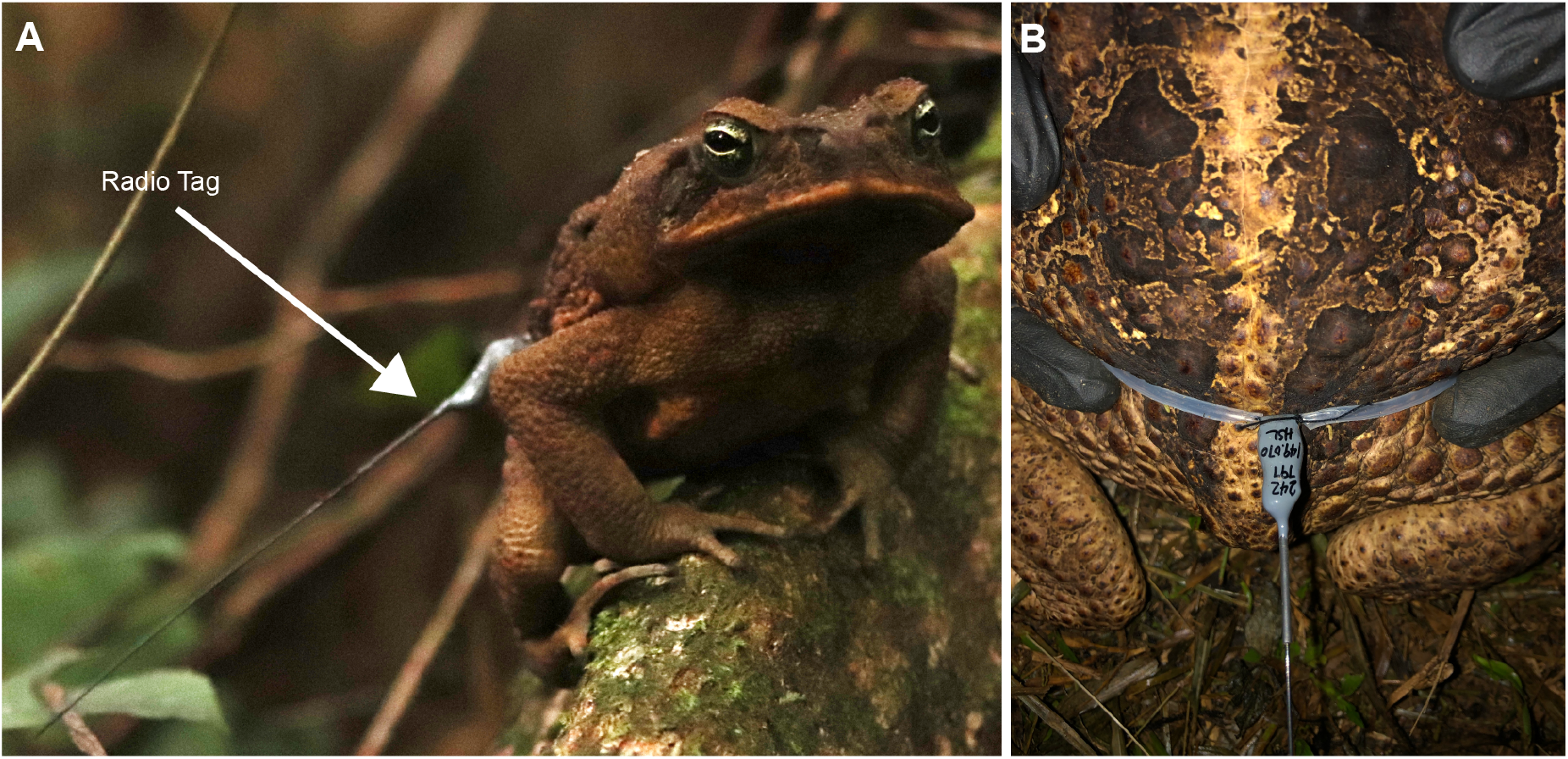
Toads tagged with radio transmitters. (A) A male toad with an attached radio transmitter. (B) Close up of radio transmitter attached to toad waist with a belt of silicone tubing.

### Tagging and baseline tracking

A radio tag (BD2, Holohil Systems Ltd, Carp, ON, Canada) was secured to a piece of 2 - 4 mm silicone tubing (Dow Corning, Midland, MI, USA,) and sized to sit loosely above the toad’s waist and legs but tightly enough that it would not slip off (Figure 1). The ends of the tubing were attached to form a loop around the toad using cotton thread, so that if the toad was to escape with the tag, the thread would disintegrate over time and free the toad. Total tag weight was less than 2 grams and negligible compared to toad mass. Tagged animals were released at their capture site after an average of 36.9 minutes (18-107 minutes). A GPS point was recorded on a handheld GPS device (RINO655t, Garmin Ltd., Schaffhausen, Switzerland) using the waypoint averaging feature. Two toads lost tags on 3 total occasions, and when they were recaptured, they were identified by dorsal wart and coloration patterns and retagged.

After release, most toads were relocated twice every 24 hours during both day and night with a flexible 3-element Yagi antenna (Biotrack Ltd, Wareham, Dorset, UK) and a SIKA radio receiver (Biotrack Ltd). Some individuals that were tagged for an extended time period and were only generally localized using the antenna without visual confirmation or a specific position being recorded on some days. In addition, one female was untagged for ∼3 weeks after an initial 3 weeks of tagging to recover from scratches from the waistband, and was retagged and tracked. Attempts were made to stay two meters away from the animals, which was sometimes difficult given the brushy environment and the need to visually validate that the transmitter was attached to the animal. Toad location was recorded as a new point if the GPS plotted the spot as being at least 5 m away from the previous location of the animal. Every week, animals were captured and inspected for any tagging-related injuries. In two instances a tag was removed because of a skin injury during baseline tracking. Later, one of these animals was recaptured and we confirmed that the wounds had healed.

The location of each individual included in our analysis was recorded over at least seven days (both before and after translocations); toads tracked for shorter durations were not included. Eleven toads (7 males and 4 females) were used to determine site fidelity for the time period of the study, defined here as non-unidirectional movement.

### Translocations

After animals had been tracked for at least 3 days, they were captured and translocated either 500 or 1000 meters in the late afternoon or night. Six individuals (five males and one female) were translocated 500 meters and 5 individuals (4 males and 1 female) were translocated 1000 m. Overall, 7 individuals were used in translocation, where 4 toads experienced trials of both distances. Translocations were repeated for 500 and 1000 m within the same animal, in random sequence, although evacuation during the COVID-19 pandemic prohibited this repeated sampling for all individuals. Translocation sites were somewhat constrained by trail access, but were done in different directions when repeated for the same animal. The release site was determined from the “outermost” baseline point for the individual. Release sites were estimated on a local GIS map with ArcPad 10.2 (ESRI, Redlands, CA, USA) and confirmed with a GPS device in the field. After capture, toads were disoriented by walking them around in a net cage for 30-60 minutes prior to release.

After translocation, toads were again located once during the day and once during the night in a 24 hour period. If possible, depending on the toad’s proximity to the camp, multiple points were taken during the night if the animal was moving, starting in the early evening. Locations were recorded in the same manner as had been done to determine home range.Animals were given ten days to return home. If they did not return after 10 days, they were returned home manually. Similarly, if the animal exhibited any extended periods of immobility (>72 hrs), they were captured and their waists checked for rubbing or wounds. One toad was injured and so the tag was removed and the animal was released at the home site. Animals were considered to be in their home area once they were within 100 meters of the translocation reference point based upon the movement range observed during baseline tracking. Following translocations, all animals were successfully untagged and released in their home areas.

### Data Analysis

GPS Data was uploaded into ArcGIS Pro 2 (ESRI) to visualize baseline points and homing trajectories. Temporal and spatial attributes of baseline activity and homing were calculated in R version 4.0.2 (R Foundation for Statistical Computing, Vienna, Austria) using the package “adehabitatLT” (Calenge, 2020a). Baseline range (maximum distance between two baseline points) was calculated by visually selecting and exporting outermost baseline points in ArcGIS and calculating distance in R. Total movement (cumulative path observed) was calculated by creating trajectory objects for individuals in adehabitatLT. The package “adehabitatHR” (Calenge, 2020b) was used to calculate Minimum Convex Polygons (MCPs) for baseline data. Coordinates of the mean center of baseline activity were calculated in ArcGIS. To account for the variability in the baseline tracking durations, we calculated the movement per day of observation as well as the proportion of days with larger (> 10 m) movements. For these calculations, any period of time greater than 48 hours in which toad location was not recorded was discarded. Straightness of homing trajectories was calculated by dividing the distance between the translocation release site and the point at which the toad was considered to be home by the cumulative movement measured. A straightness index of 1 indicates a toad that travelled in a completely straight line, with values approaching 0 indicating less direct routes taken. Normality of data was determined by Shapiro-Wilks Test. Comparisons for normal data were performed with a t-test, and comparisons of non-parametric data were performed with a Mann-Whitney U Test. All statistical tests of significance were performed in R (version 4.0.2).

### Permits and Ethical Statement

The experiments were conducted in strict accordance with the European, French and USA laws and following the ‘Guidelines for use of live amphibians and reptiles in the field and laboratory research’ by the Herpetological Animal Care and Use Committee (HACC) of the American Society of Ichthyologists and Herpetologists (Beaupre et al., 2004) and the Association for the Study of Animal Behaviour (ASAB) “Guidelines for the treatment of animals in behavioral research and teaching” (Vitale et al., 2018). This experiment was performed under approval from the scientific committee of the Nouragues Ecological Research Station. All procedures were approved by the Institutional Animal Care and Use Committee of Stanford University (protocol #33714).

## Results

### Baseline Tracking

Ten toads out of eleven exhibited site fidelity during 7-55 tracking days (mean = 24 days, s.d. = 15 days). Adjusting for periods of at least 48 hours in which observations of position were not made, space use was quantified over 7-32 days (mean = 18 days, s.d. = 10 days). The observed range of the ten toads (Figure 2A) varied from 26.8 m to 236.7 meters (mean = 109.6 meters, s.d. = 73.9), which we interpret as site fidelity. Our statistical tests revealed no sex difference in baseline parameters, although the sample size was not large enough to detect a sex difference. On average, toads moved more than 10 m on 60% of days of observation, with many periods lacking substantial movement.

**Figure 2.**
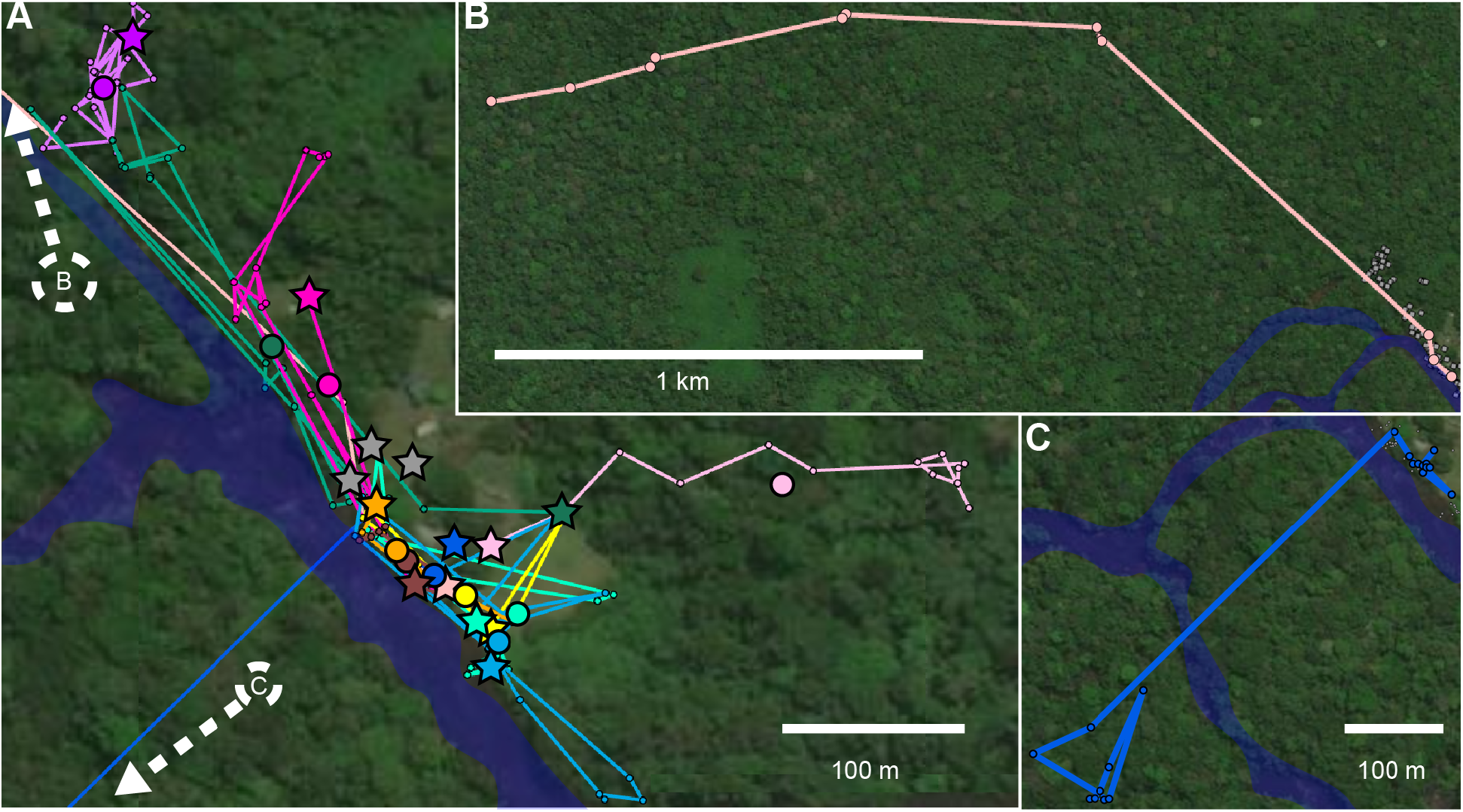
Baseline tracking of *Rhinella marina*. (A) Baseline tracking for 10 toads (over 7-55 days) showing site fidelity; colors represent individuals. Location points are connected in temporal order. Larger circles represent the mean center of baseline position. Stars represent tagging locations and gray stars represent toads that were tagged but not substantially tracked. (B) A female toad moved 2 kilometers without translocation. (C) A male toad was initially localized near other males, but was moved across the river while in amplexus.

During baseline tracking, males were observed more often and in higher density than females. A minimum convex polygon encapsulating the mean centers of the for seven males exhibiting site fidelity measured 858.6 m^2^. In contrast, an MCP for the mean centers of the three females exhibiting site fidelity measured 8328.8 m^2^. Many other males were observed within the same vicinity of the tracked males.

One female toad not exhibiting site fidelity was tracked for 16 days and moved a cumulative distance of 1999.4 m for a 1779.2 m displacement (Figure 2B). Another male toad exhibited site fidelity until it was displaced more than 300 meters across the river to an island by an untagged female while in amplexus (Figure 2C). The male was first observed in amplexus in its home range, and relocated on the other side of the river four days later. The toad was untagged after 15 days on the island (of which it was in amplexus for 6), and did not return to its previous location for the duration of the observation.

### Translocations and Homing

Of the six individuals translocated 500 meters, five returned to their home areas within three days (Figure 3A). Returns were observed between 27.5 and 79.9 hours after release at the translocation site (mean = 57.5 hours, s.d. = 21.6 hours) (Figure 3B). Observed straightness of four homing paths with en-route relocations varied from 0.89 to 0.98 (mean = 0.94, s.d. = 0.04), indicating that the toads returned directly with minimal exploration. One individual did not return home, traveling 250 meters in the wrong direction before returning to its translocation site. This toad was manually returned home because of observed abrasions from the belt following 2 days of no movement.

**Figure 3.**
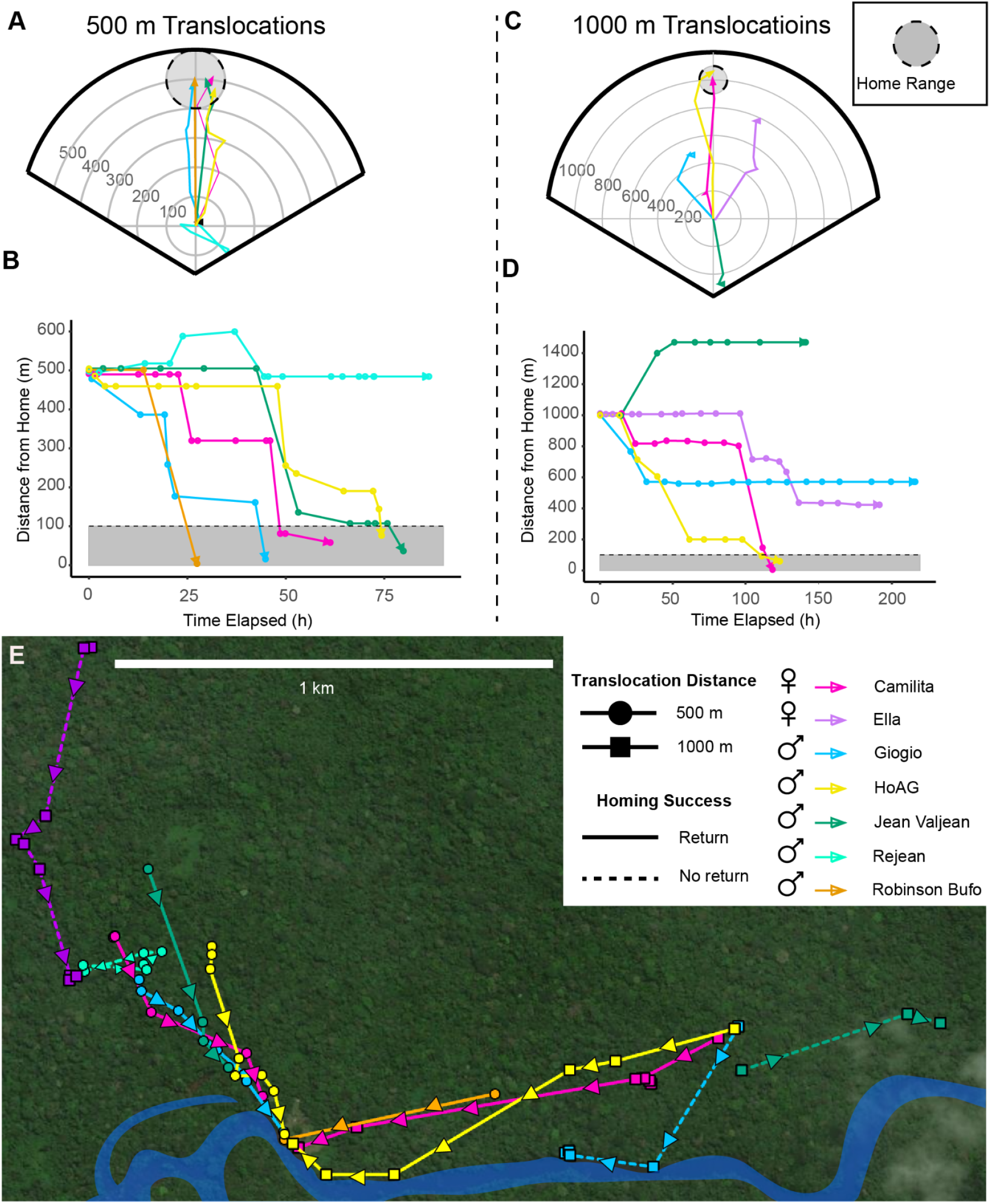
Homing trajectories of translocated *Rhinella marina*. (A) Homing trajectories of toads translocated 500 meters. Plot center represents translocation sites and gray circle represents home area. (B) Distance from home plotted against time after translocation during homing from 500 m, with a gray rectangle representing the home area. (C) Homing trajectories and (D) distance from home over time for toads translocated 1000 m. (E) Movement of toads after translocation on a map of the field site; solid lines show movement following 500 meter translocations and dashed lines represent 1000 meter translocations.

Of the five individuals translocated 1000 m, two returned to home sites within ∼5 days (Figure 3C). Returns took 123.7 and 118.7 hours (Figure 3D). Both returned directly with measured straightness of 0.94. Both animals were observed to make their first substantial movements after 24 hours post-translocation (24.25 and 26.67 hours), and both exhibited secondary periods of immobility following this initial movement. Two of the toads that did not return home initially moved in the correct direction before stopping prior to reaching their home areas. The remaining individual moved ∼500 m in the wrong direction before becoming stationary.

## Discussion

The study of navigation has focused on animals that move large distances or perform tasks requiring fine scale navigation. This may leave the abilities of other species capable of precise navigation overlooked. We showed for the first time that cane toads are able to directly navigate back to home sites following translocations of up to one kilometer.

Using radio-tracking of toad movements prior to translocation, we found that toads were largely sedentary without translocation and movements were largely nocturnal, similar to previous observations in both native and invasive cane toads, (Carpenter and Gillingham, 1987; DeVore et al., 2021; Ward-Fear et al., 2016; Zug and Zug, 1979). When compared to the recent tracking of toads done in French Guiana (DeVore et al., 2021), the minimal shifts in position in our toads parallel the activity of toads tracked in the rainforest as compared to other habitats. We did not observe larger scale foraging movements (<100 meters) that have been reported in coastal toads, although the movement of a female toad for almost 2 kilometers greatly exceeds any movements reported in coastal or rainforest sites. Overall, our observations here generally align with previous studies, although more research is needed to gain a better understanding of movement ecology and how it covaries with biotic and abiotic factors.

During our tracking, males were clustered in relatively high density by the river, with females spread out in the surrounding area. These observations suggest female cane toads may move to areas populated by males to mate and return back to their original home areas, but more detailed long-term tracking studies are needed to understand sex specific spatial distribution and movements. Studies of invasive toads in Hawaii have shown some sex differences in space use, but not necessarily with respect to range and total inhabited area (Ward-Fear et al., 2016). Tracking of native range toads showed that the only variable affected by sex was the probability of emerging to forage, with males more lilkely to emerge (Devore et. al, 2021), although we did not consider this parameter. Our observations of large scale movements are comparable to those recorded in invasive toads (Schwarzkopf and Alford, 2002; Ward-Fear et al., 2016), but similar observations have not been reported in native-range cane toads. In Australia, the available range for expansion may contribute to their propensity to disperse (Brown et al., 2014; Phillips et al., 2006). A longer term space use study is necessary to better characterize full range and complexity of cane toad movement in their native habitat.

Toads showed the ability to directly navigate back to home sites following translocations of 500 and 1000 m. Previous studies have indicated that cane toads are good navigators, but this ability has not been quantified in detail. A previous homing study in Panama showed that cane toads could return to specific lights to hunt for insects following short distance translocations of <100 meters (Brattstrom, 1962). In Australia, some cane toads are able to return from over a kilometer to bird ground nests they had exploited for food (Boland, 2004). Despite a very different life history and nocturnal activity, cane toads showed similar navigational accuracy to poison frogs homing in the same habitat from lesser distance (Pašukonis et al., 2014b) and straighter trajectories than poison frogs moving comparable distances (Pašukonis et al., 2018), although this may be partially attributable to the lower resolution of toad trajectories. This straightness of the homing behavior contrasts with the general meandering of regular movements made by native range toads (DeVore et al., 2021). Similar to poison frogs, toads were initially stationary at their translocation sites, suggesting a period of gathering bearings (Pašukonis et al., 2014a; Pašukonis et al., 2014b; Pašukonis et al., 2018). Even toads that did not home successfully showed the ability to orient, with two of the three toads that did not return from 1000 meters moving in the correct direction before stopping short of their observed home range. The season at the field site was unusually dry and it did not rain for the duration of these translocations. Cane toads have different spatial behaviors based upon rainfall and the availability of water resources (DeVore et al., 2021), and the failure to home in our study may potentially have been due to stopping at available water resources. Overall, toads demonstrated a capacity to return with accuracy to their home areas from distances exceeding their regular movements.

The actual sensory mechanisms that allow for this homing are unknown. Magnetoreception and olfaction have been shown to play an important role in navigation in other bufonid species in smaller scale translocations (Sinsch, 1990; Sinsch, 1987; Sinsch, 1992). It is possible that, since males were largely congregated in one area, that auditory taxis to male calling could have contributed to homing. However, males were not calling throughout the entire study and it is unlikely that calls would travel a kilometer through dense forest. The use of visual cues is unlikely, given that toads did not explore the area and the dense rainforest understory results in many obstacles. Simple olfactory taxis to water also seems unlikely in the case of successfully returning animals, given that toads could have returned to stretches of river closer to their translocation sites (Figure 3E). Multiple navigation mechanisms (beaconing, path integration, etc.) have been described in amphibians (Sinsch, 2006) and more work should be done to identify which (if any) of these paradigms are applicable to the cane toad’s apparent ability to navigate.

Our observations show that cane toads are capable of advanced navigation over long distances after displacement from a home area. It is thus possible that well-developed navigational abilities are ancestral and widely shared among amphibians. Cane toads are common, large and invasive amphibians able to carry additional biologging devices (like accelerometers and GPS), making them particularly interesting for field and lab studies on amphibian navigation. Future research could include testing toad navigation under conditions where senses are manipulated to identify which sensory cues are important for navigation, as well as identifying the neural basis of navigation in amphibians.

## Supporting information

Baseline Tracking Data

Translocation Data

## Acknowledgements

We would like to thank the technical team of the Nouragues Ecological Research Station for their support: Wémo Betian, Patrick Chatelet, Elodie Courtois, Philippe Gaucher, Florian Jeanne, Bran Leplat, and Nina Marchand. We would also like to thank Agaci Dutra de Souza for transportation of personnel and equipment, and Charles Albert Tropée for piloting our COVID-19 evacuation flight. We are grateful to the staff of the Nouragues Nature Reserve for their commitment to preserving our natural world. This work is part of a partnership between AP and the Nouragues Nature Reserve aimed at improving and spreading the knowledge about amphibians. We thank Marie-Therese Fischer and Matthew Greenlees for reading this paper and providing invaluable comments. We acknowledge that the field work was done in a region including the unceded and ancestral lands of native peoples of the Guiana Shield, who were displaced by force and disease from European colonists. We acknowledge that Stanford University resides on the unceded and ancestral lands of the Muwekma Ohlone people.

## Author contributions

Conceptualization: DAS (equal), AP (equal)

Methodology: DAS (equal), AP (equal)

Formal Analysis: DAS

Investigation: DAS

Resources: LAO (primary), AP (supporting)

Data Curation: DAS

Writing – original draft preparation: DAS

Writing – review and editing: DAS (equal), LAO (equal), AP (equal)

Visualization: DAS

Supervision: LAO (primary), AP (supporting)

Project administration: LAO (primary), AP (supporting)

Funding acquisition: DAS (supporting), LAO (primary), AP (supporting)

## Data Accessibility

Spreadsheets with raw data for baseline and translocation information are uploaded with supplementary information.

